# Antiretroviral treatment reveals a novel role for lysosomes in oligodendrocyte maturation

**DOI:** 10.1101/2022.08.05.502855

**Authors:** Lindsay K. Festa, Abigail E. Clyde, Caela C. Long, Lindsay M. Roth, Judith B. Grinspan, Kelly L. Jordan-Sciutto

**Author notes:** Correspondence should be addressed to: Kelly L. Jordan-Sciutto, Department of Oral Medicine, School of Dental Medicine, University of Pennsylvania, 240 S. 40^th^ St, Philadelphia, PA 19104., Judith B. Grinspan, Department of Neurology, The Children’s Hospital of Philadelphia, 516D Abramson Center, 3615 Civic Center Blvd, Philadelphia, PA 19104.

## Abstract

White matter deficits are a common neuropathologic finding in neurologic disorders, including HIV-associated neurocognitive disorders (HAND). In HAND, the persistence of white matter alterations despite suppressive antiretroviral (ARV) therapy suggests that ARVs may be directly contributing to these impairments. Here, we report that a frontline ARV, bictegravir (BIC), significantly attenuates remyelination following cuprizone-mediated demyelination, a model that recapitulates acute demyelination, but has no impact on already formed mature myelin. Mechanistic studies in vitro revealed that treatment with BIC leads to significant decrease in mature oligodendrocytes accompanied by lysosomal de-acidification and impairment of lysosomal degradative capacity with no alterations in lysosomal membrane permeability or total lysosome number. Activation of the endolysosomal cation channel TRPML1 prevents both lysosomal de-acidification and impairment of oligodendrocyte differentiation by BIC. Lastly, we show that de-acidification of lysosomes by compounds that raise lysosomal pH is sufficient to prevent maturation of oligodendrocytes. Overall, this study has uncovered a critical role for lysosomal acidification in modulating oligodendrocyte function and has implications for neurologic diseases characterized by lysosomal dysfunction and white matter abnormalities.

**Table of Contents:** *Main Points:*

- The antiretroviral, bictegravir, inhibited remyelination through OPC differentiation blockade and had no effect on mature myelin
- Bictegravir inhibits oligodendrocyte differentiation through de-acidification of lysosomes and this was prevented via activation of the lysosomal channel TRPML1
- De-acidification of lysosomes by other drugs (e.g. bafilomycin A) is sufficient to inhibit oligodendrocyte maturation

**Table of Contents Image:** 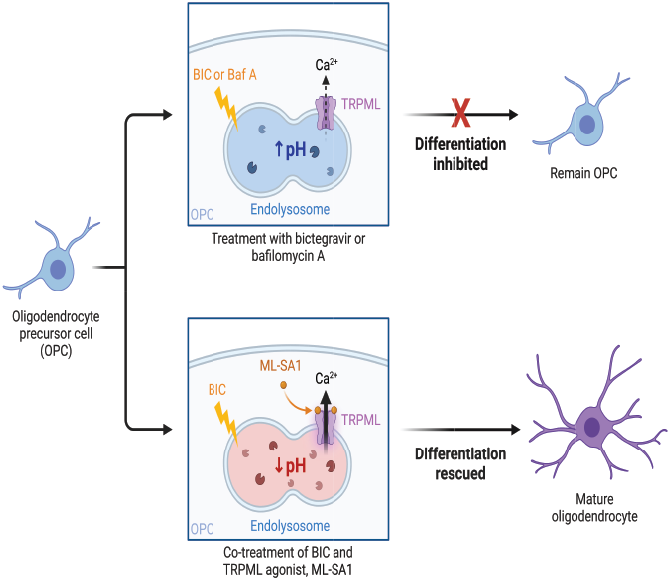

## Introduction

In the central nervous system (CNS), oligodendrocytes form myelin, the concentrically wrapped membrane that insulates axons and facilitates rapid and efficient action potential conduction (Chapman and Hill, 2020). Oligodendrocytes arise from a self-renewing pool of oligodendrocyte precursor cells (OPCs), which populate the entire CNS and undergo a burst of differentiation and subsequent myelination in development, followed by ongoing, but slow, OPC differentiation and myelination throughout life (Hill et al., 2018; Hughes et al., 2018). In addition to allowing saltatory conduction along axons, oligodendrocytes perform numerous other functions in the developing and adult CNS including, but not limited to, internode maturation and maintenance, regulation of neuronal excitability, and metabolic support of axons (Pepper et al., 2018). Based on these functions, as well as their direct physical contact with neurons, oligodendrocytes are well-positioned to exert significant influence over neuronal health and circuitry; this is further highlighted by white matter disorders, such as multiple sclerosis, that result in motor, cognitive, and behavioral impairment (Filippi et al., 2018). While extensive research has been conducted on the motility, proliferation, and differentiation of these cells, there are still outstanding questions regarding how these processes are regulated.

The lysosome is a membrane-enclosed vesicular organelle that contains more than 60 acidic hydrolases, which break down cell components and complex macromolecules into the constituent building blocks (Ballabio and Bonifacino, 2020). Based on their initial characterization, these organelles were long regarded as the “garbage-disposal” of the cell; however, emerging evidence has demonstrated that lysosomes regulate diverse processes, including nutrient sensing, metabolic regulation, gene regulation, and plasma membrane repair (Bonam et al., 2019). In addition to general homeostatic cell processes, lysosomes regulate cell-type specific functions including in oligodendrocytes. Perhaps the most well-defined role of lysosomes in oligodendrocytes is the release of proteolipid protein (PLP), a major component of myelin, from late endosomes/lysosomes to the plasma membrane. PLP is expressed in the rough endoplasmic reticulum where it then undergoes transport to the Golgi and eventually the plasma membrane following signals from the neuron (Trajkovic et al., 2006). In addition to PLP, endocytic sorting and recycling has been observed for two other myelin proteins, myelin associated glycoprotein (MAG) and myelin oligodendrocyte glycoprotein (MOG), as well as the OPC proteoglycan NG2 (Winterstein et al., 2008; Daynac et al., 2018). Lastly, recent work has demonstrated that autophagy, a catabolic function that is dependent on the lysosome, is responsible for proper myelin compaction possibly via clearance of cytoplasm (Bankston et al., 2019). The low intraluminal pH of the lysosome is critical for these diverse functions as lysosomal dysfunction of neurons is observed in many neurodegenerative diseases (Bonam et al., 2019).

Lysosomal acidification defects have been observed in neurons of several CNS disorders, including Alzheimer’s disease, Parkinson’s disease, multiple system atrophy, and HIV-associated neurocognitive disorders (HAND) (Kreher et al., 2021). Despite effective viral suppression from combination antiretroviral therapy (cART), approximately 50% of people living with HIV (PWH) present with cognitive, motor, and behavioral deficits collectively known as HAND (Saylor et al., 2016). White matter dysfunction, including thinning of the corpus callosum, loss of structural integrity, and dysregulation of transcripts critical for myelination, have persisted despite cART adherence, suggesting that antiretrovirals (ARVs) themselves may contribute to white matter injury and HAND (Borjabad et al., 2011; Tate et al., 2011; Solomon et al., 2019). Previous work in the laboratory reported that ARV compounds from different classes, such as protease inhibitors (darunavir and saquinavir) and integrase strand transfer inhibitors (INSTI; elvitegravir), inhibit oligodendrocyte maturation *in vitro* via lysosomal deacidification and the integrated stress response (Festa et al., 2021; Roth et al., 2021).

Here we report that frontline INSTI, bictegravir (BIC), inhibits remyelination, a process dependent on oligodendrocyte differentiation, following cuprizone-mediated demyelination. Our mechanistic studies in vitro demonstrate that lysosomal de-acidification mediates BIC-induced inhibition of oligodendrocyte maturation and can be rescued via lysosomal calcium efflux. Lastly, we show that de-acidification of lysosomes by compounds that raise lysosomal pH is sufficient to prevent maturation of oligodendrocytes. Overall, this study has uncovered the critical role of lysosomal acidification in modulating oligodendrocyte differentiation and has implications not only for HAND but other neurologic diseases characterized by lysosomal dysfunction and white matter abnormalities.

## Materials and Methods

### Reagents

Reagents and chemicals used were LysoTracker Red DND-99 (ThermoFisher Scientific; L7528), LysoSensor Yellow/Blue DND-160 (ThermoFisher Scientific; L7545), DAPI (ThermoFisher Scientific; D1306), DQ-BSA Red (ThermoFisher Scientific; D12051), FluoroMyelin Green (ThermoFisher Scientific; F34651), bictegravir (Toronto Research Chemicals; B382095), bafilomycin A (Millipore Sigma; B1793), Leu-Leu methyl ester hydrobromide (Millipore Sigma; L7393), MLSA1 (Millipore Sigma; SML0627), and bis-cyclohexanone-oxaldihydrazone (cuprizone; Sigma Aldrich; C9012).

### Antibodies

The following antibodies were used for immunohistochemical analysis of brain tissue sections: aspartoacylase (ASPA; mature oligodendrocytes; GeneTex; GTX110699; 1:500), neural/glial antigen 2 (NG2; OPCs; Millipore; AB5320; 1:200), ionized calcium binding adaptor molecule 1 (Iba1; microglia; Wako; 019-19741; 1:500), and glial fibrillary acidic protein (GFAP; astrocytes; hybridoma supernatant courtesy of Dr. Virginia Lee, University of Pennsylvania, 1:2). The following antibodies were used for *in vitro* immunocytochemistry: A2B5 (OPCs; mouse IgM hybridoma; 1:1) (Eisenbarth et al., 1979), galactocerebroside (GalC; immature oligodendrocytes; mouse IgG3 hybridoma; 1:5) (Raff et al., 1978), proteolipid protein (PLP; mature oligodendrocytes; AA3 rat hybridoma; 1:1), galectin 3 (Abcam; ab2785; 1:100), and LAMP1 (Enzo Life Sciences; ADI-VAM-EN001-F; 1:100). The following antibodies were utilized for immunoblotting: myelin basic protein (MBP; Biolegend; 808401; 1:1000), PLP (rat hybridoma, 1:1000), and alpha tubulin (α-tubulin; Sigma-Aldrich; T5168; 1:10,000).

### Animal Use

All experiments were conducted in accordance with the guidelines set forth by the Children’s Hospital of Philadelphia Institutional Animal Care and Use Committee.

### Mass spectrometry study

C57BL/6 female mice 15-16 weeks old (Jackson Laboratories, PA) were used to determine the maximum plasma concentration of BIC via mass spectrometry. Fifteen mice underwent oral gavage and were treated with one of three doses of BIC: 2 mg/kg, 4 mg/kg, or 8 mg/kg. Blood was collected via the submandibular vein after 3 hrs and sera was isolated and processed for mass spectrometry analysis. Based on the data, animals were treated with 8 mg/kg BIC following cuprizone-mediated demyelination (see below).

### Cuprizone model and *in vivo* drug treatments

10-12 weeks old female C57BL/6 mice were used for the cuprizone demyelination model and were divided into two groups: control (powdered chow) and cuprizone in powdered chow (0.2% w/v). Each group was further subdivided into untreated, vehicle (5% DMSO), or BIC-treated (8 mg/kg diluted in 5% DMSO), with five mice per group for a total of 40 animals. After 5 weeks of either control or cuprizone feeding, five control mice and five cuprizone mice were perfused and their brains were processed for immunohistochemistry (see below). The other mice underwent once daily oral gavage with either DMSO or BIC (except for the untreated group) while being fed conventional diet during the last 3 weeks of the experiment (recovery period). At the end of the recovery period, all animals were sacrificed and their brains were processed for immunohistochemistry.

### Immunohistochemistry

Mice were anesthetized with a ketamine/xylazine cocktail and were intracardially perfused first with ice-cold PBS and then 4% paraformaldehyde (pH 7.4). Brains were removed and the olfactory bulb and cerebellum were removed using a brain matrix. Brains were postfixed in 4% paraformaldehyde for 24 hours followed by immersion in 30% sucrose solution for at least 24 hrs. Brains were cryopreserved and embedded in optimal cutting temperature compound (OCT) followed by coronal sectioning (12 μm) on a cryostat (Leica Microsystems, Exton, PA). The third ventricle was used as a regional anatomical marker for the region of the corpus callosum most consistently demyelinated by cuprizone. Sections were blocked in 10% normal goat serum for 1 hr at RT and then incubated overnight with primary antibodies diluted in 2% normal goat serum in PBS. The next day, sections were rinsed three times in 1X PBS and incubated with fluorescent secondary antibodies in 1X PBS (1:200) for 1 hr at RT. Sections were rinsed three times in 1X PBS prior to being counterstained with DAPI for 5 mins and mounted in ProLong Gold anti-fade reagent (ThermoFisher Scientific; P36930). For FluoroMyelin Green staining, sections were incubated for 20 minutes in 1X PBS (1:200) followed by counterstaining with DAPI and mounting in ProLong Gold. Images were acquired on a DMi8 Leica inverted confocal microscope (Leica Microsystems) using a 40x objective/1.3 NA. To quantify mean integrated density (NG2, FluoroMyelin, GFAP, and Iba1), five 150 mm x 150 mm regions of interest from the corpus callosum across three sections from each animal were analyzed. The channel of interest was separated and thresholded to isolate staining from background, and integrated density was calculated using ImageJ. Quantification of cell number (ASPA) was performed in a similar manner and the number of ASPA+ cells were hand counted using ImageJ.

### Primary rat oligodendrocyte precursor cell cultures

Primary rat glial cells were isolated from brains of postnatal day 1 Sprague Dawley rats (Charles River Laboratories, Wilmington, MA) and plated on T75 flasks (Jensen et al., 2015). Upon reaching confluency, OPCs were purified using the “shake-off” method (McCarthy and de Vellis, 1980). Briefly, T75 flasks were rotated on an orbital shaker set to 250 rpm and incubated overnight at 37°C. The following day, cells were filtered using a 20 μm nylon net (Merck Millipore, Darmstadt, Germany), followed by centrifugation at 1500 rpm for 5 mins at 4°C. The supernatant was discarded, and the pellet was resuspended in 5 mL Neurobasal medium supplemented with B27 and incubated on a bacteriological petri dish for 15 mins at 37°C. The supernatant was collected and centrifuged at 1500 rpm for 5 mins at 4°C. The pellet was resuspended in Neurobasal medium supplemented with B27 and growth factors: PDGF (2 ng/mL), NT3 (1 ng/mL), and FGF (10 ng/mL) and plated on 24-well plates with coverslips (immunocytochemistry), 10 cm dishes (immunoblotting), or 96-well plate (lysosomal pH measurements).

### Drug treatments

Primary rat OPCs were grown in 24-well plates with coverslips or 10 cm dishes, for immunocytochemistry or immunoblotting respectively, until they reached approximately 70% confluency. In order to differentiate OPCs into mature oligodendrocytes, growth medium was replaced with differentiation medium containing 50% DMEM, 50% Ham’s F12, pen/strep, 2 mM glutamine, 50 μg/mL transferrin, 5 μg/mL putrescine, 3 ng/mL progesterone, 2.6 ng/mL selenium, 12.5 μg/mL insulin, 0.5 μg/mL T4, 0.3% glucose, and 10 ng/mL biotin. At the time of differentiation, cells were treated with vehicle (DMSO), bictegravir (1.35 μM, 13.5 μM, or 41 μM), bafilomycin A (250 pM, 500 pM, or 750 pM), or MLSA1 (20 μM) for 72 hrs prior to staining or protein collection. Leu-Leu-OMe (1 mM) was given 2 hrs prior to experimental end points. On average, typically 30-50% of OPC differentiation into mature oligodendrocytes is observed in untreated conditions.

### Immunoblotting

Whole cell extracts of primary rat oligodendrocytes were prepared in ice cold lysis buffer (25 mM Tris (pH 7.4), 10 mM EDTA, 10% SDS, 1% Triton-X 100, 150 mM NaCl, protease and phosphatase inhibitor cocktail) followed by sonication and centrifugation at 10,000 rpm at 4°C for 30 mins. Protein concentrations were by spectrophotometer ND-1000 (ThermoFisher Scientific). Protein (7.5 μg) were loaded onto a 4-12% Bis-Tris gradient gel and after separation, proteins were transferred to Immobilin-FL membranes. Membranes were blocked in 5% milk for 1 hour at RT and then incubated overnight at 4°C with primary antibodies in 5% BSA. Following three washes in TBS-T, membranes with incubated with fluorescent probe-conjugated secondary antibodies in 5% BSA. Membranes were visualized using an Odyssey Infrared Imaging System (LiCOR). Densitometric analysis of band intensities were conducted using ImageJ. All bands were normalized to the loading control.

### Immunocytochemistry

Following drug treatments, coverslips were washed three times in DMEM/F12 and incubated in primary antibodies to surface antigens (A2B5 and/or GalC) for 30 mins at RT. Coverslips were then rinsed three times with DMEM/F12 and incubated in appropriate fluorescent conjugated secondary antibodies for 30 mins at RT. Cells were then rinsed three times in DMEM/F12, fixed in 4% paraformaldehyde for 10 mins, and permeabilized with 0.1% Triton-X 100 for 5 mins. Cells were then incubated in primary antibodies to internal antigens (PLP, LAMP1, and/or Gal3) for 30 mins at RT. Coverslips were rinsed three times in 1X PBS and incubated in appropriate secondary antibodies for 30 mins at RT. Lastly, cells were rinsed, counterstained with DAPI for 5 mins, and then mounted on slides with ProLong Gold anti-fade reagent. For propidium iodide staining, the same steps were followed with the omission of permeabilization. Cells were imaged either using a Keyence BZ-X-700 digital fluorescent microscope (Keyence Corporation) for OPC and oligodendrocyte cell counts or a DMi8 Leica inverted confocal microscope equipped with a 40x objective/1.3 NA. Quantification of oligodendrocyte lineage counts were conducted by hand counting the number of OPCs, immature oligodendrocytes, mature oligodendrocytes, and DAPI+ cells across 25 fields/coverslips and 3 coverslips per condition for each biological replicate. The percentage of each cell type was calculated and normalized to the untreated condition. For LAMP1 and Gal3 puncta counts, imaging fields were chosen at random and Z-stacks from 6 fields/coverslips and 3 coverslips/condition for each biological replicate were acquired. The channel of interest was separated and thresholded to isolate the puncta (0.5 μm^2^-1.0 μm^2^). The number of puncta per cell were quantified using the particle analysis plugin in ImageJ.

### TUNEL assay

Cells were fixed with ice cold methanol for 10 mins, washed with PBS, and permeabilized with 0.1% Triton-X 100 and 0.5% BSA in PBS for 5 mins. Positive control coverslips were generated during this time by incubating in DN buffer (30 mM Tris base (pH 7.2), 140 mM sodium cacodylate, 4 mM magnesium chloride, and 0.1 mM dithiothreitol) for 2 mins, followed by DNase (1:200) in DN buffer for 10 mins. All coverslips were washed with PBS and then placed in TDT buffer (30 mM Tris base (pH 7.2), 140 mM sodium cacodylate, 1 mM cobalt chloride) for 2 mins. Cells were incubated for 1 hr at 37°C with TdT and biotin-UTP in TDT buffer. After subsequent PBS washes, cells were placed in TB buffer (300 mM sodium chloride, 30 mM sodium citrate) for 15 mins and then 2% BSA solution for 30 mins. Finally, cells were incubated with rhodamine-conjugated streptavidin for 20 mins, rinsed in PBS, incubated with DAPI for 5 mins, and then mounted on slides with ProLong Gold. Images were acquired using a Keyence BZ-X-700 fluorescent microscope using a 40x/0.7 NA objective.

### LysoSensor Yellow/Blue experiments

After 72 hrs of differentiation and drug treatments, differentiation medium was removed and fresh media with 5 μM LysoSensor Yellow/Blue was added for 5 mins. Cells were then rinsed three times in isotonic solution (105 mM sodium chloride, 5 mM potassium chloride, 10 mM HEPES buffer, 5 mM sodium bicarbonate, 60 mM mannitol, 5 mM glucose, 0.5 mM magnesium chloride, and 1.3 mM calcium chloride) and then left in isotonic solution for imaging. For generation of the standard curve, pH standards (4.0-6.5) were generated by preparing a 10 μM H^+^/Na^+^ ionophore monensin and 20 μM H^+^/K^+^ ionophore nigericin dissolved in 20 mM 2-(N-morpholino)ethane sulfonic acid (MES), 110 mM potassium chloride, and 20 mM sodium chloride followed by adjustment of pH with hydrogen chloride and sodium hydroxide. Data was generated by exciting each well at 340 nm and 380 nm with emission at 520 and the ratio between each excitation was calculated followed by extrapolation based on the standard curve.

### LysoTracker Red experiments

Following 72 hrs of drug treatments, cells were incubated with 50 nM LysoTracker Red DND-99 in fresh differentiation medium for 30 mins at 37°C. After this incubation, cells were rinsed three times with DMEM/F12 and immunocytochemistry for oligodendrocyte lineage markers was performed as described above. Images were acquired on a DMi8 Leica inverted confocal microscope equipped with a 40x/1.3 NA objective and 3x optical multiplier (120x total magnification). Imaging fields were chosen at random and Z-stacks from 6 fields/coverslips and 3 coverslips/condition for each biological replicate were acquired. The channel of interest was separated and thresholded to isolate the puncta (0.5 μm^2^-1.0 μm^2^). The number of puncta per cell were quantified using the particle analysis plugin in ImageJ.

### DQ-BSA Red experiments

Differentiated oligodendrocytes were treated with 10 μg/mL in fresh differentiation medium for 1 hr or 6 hrs at 37°C. Cells were then rinsed three times with DMEM/F12 and then stained for oligodendrocyte lineage markers as outlined above. Images were acquired on a DMi8 Leica inverted confocal microscope equipped with a 40x/1.3 NA objective and 3x optical multiplier (120x total magnification). Imaging fields were chosen at random and Z-stacks from 6 fields/coverslips and 3 coverslips/condition for each biological replicate were acquired. The channel of interest was separated and thresholded to isolate the puncta (0.5 μm^2^-1.0 μm^2^). The number of puncta per cell were quantified using the particle analysis plugin in ImageJ.

### Experimental design and statistical analysis

The number of animals used for *in vivo* studies was calculated by power analysis using G* Power version 3.1.9.4 (Faul et al., 2009). Sample sizes were calculated using an independent measures study design with α= 0.05 and power= 0.8. Female C57BL/6 mice were randomly assigned to groups for the cuprizone-mediated demyelination experiments. All *in vitro* experiments were performed with at least N= 3 biological replicates, and each biological replicate used OPCs derived from a different litter of P0-P4 rat pups. Outliers were neither encountered nor removed for any analysis. Data are represented as mean ± SEM for *in vitro* and *in vivo* experiments. All *in vivo* and *in vitro* experiments, except for those with MLSA1, were analyzed using a one-way ANOVA followed by Tukey post hoc. Experiments involving MLSA1 were assessed using a two-way ANOVA followed by Tukey’s multiple comparisons test. A p<0.05 was considered statistically significant. P and n are reported in the figure legends. All statistical analysis was performed with GraphPad Prism, version 9.0 (GraphPad Software).

## Results

### Bictegravir significantly inhibits remyelination following cuprizone-mediated demyelination

We have previously shown that the INSTI elvitegravir blocks remyelination following a demyelinating insult; thus, we sought to investigate whether another INSTI, bictegravir, exhibited similar deleterious effects (Roth et al., 2021). Remyelination is the repair process by which myelin is regenerated, which is driven primarily by the differentiation of OPCs into mature, myelinating oligodendrocytes. In order to model robust demyelination and remyelination, we utilized a widely employed toxin-mediated demyelination mouse model. The copper chelator bis-cyclohexanone-oxaldihydrazone (cuprizone) delivered in the diet for 5 weeks results in widespread demyelination in the brain, including the corpus callosum, the largest white matter tract in the mouse brain (Matsushima and Morell, 2001). Switching the animals back to normal diet results in remyelination over the course of 3 weeks (the recovery period). During this recovery period, animals were administered BIC (8 mg/kg) via oral gavage once daily, while control animals received either DMSO vehicle or no gavage (Figure 1A). Additionally, animals that were fed control diet throughout the 8-week period were also treated with DMSO or BIC (8 mg/kg). This dose of BIC, determined by mass spectrometry, was comparable to physiologically relevant plasma concentrations observed in patients (Table 1; (Gallant et al., 2017). A subset of animals was sacrificed at 5 weeks to determine the extent of the demyelinating lesion.

**Table 1.** Comparison of maximum plasma concentration of bictegravir in mice versus humans.

| **Species** | **Average maximum plasma concentration (μM)** |
| --- | --- |
| Human | 13.53 μM (Gallant et al., 2017) |
| Mice | 13.88 μM |

**Figure 1.**
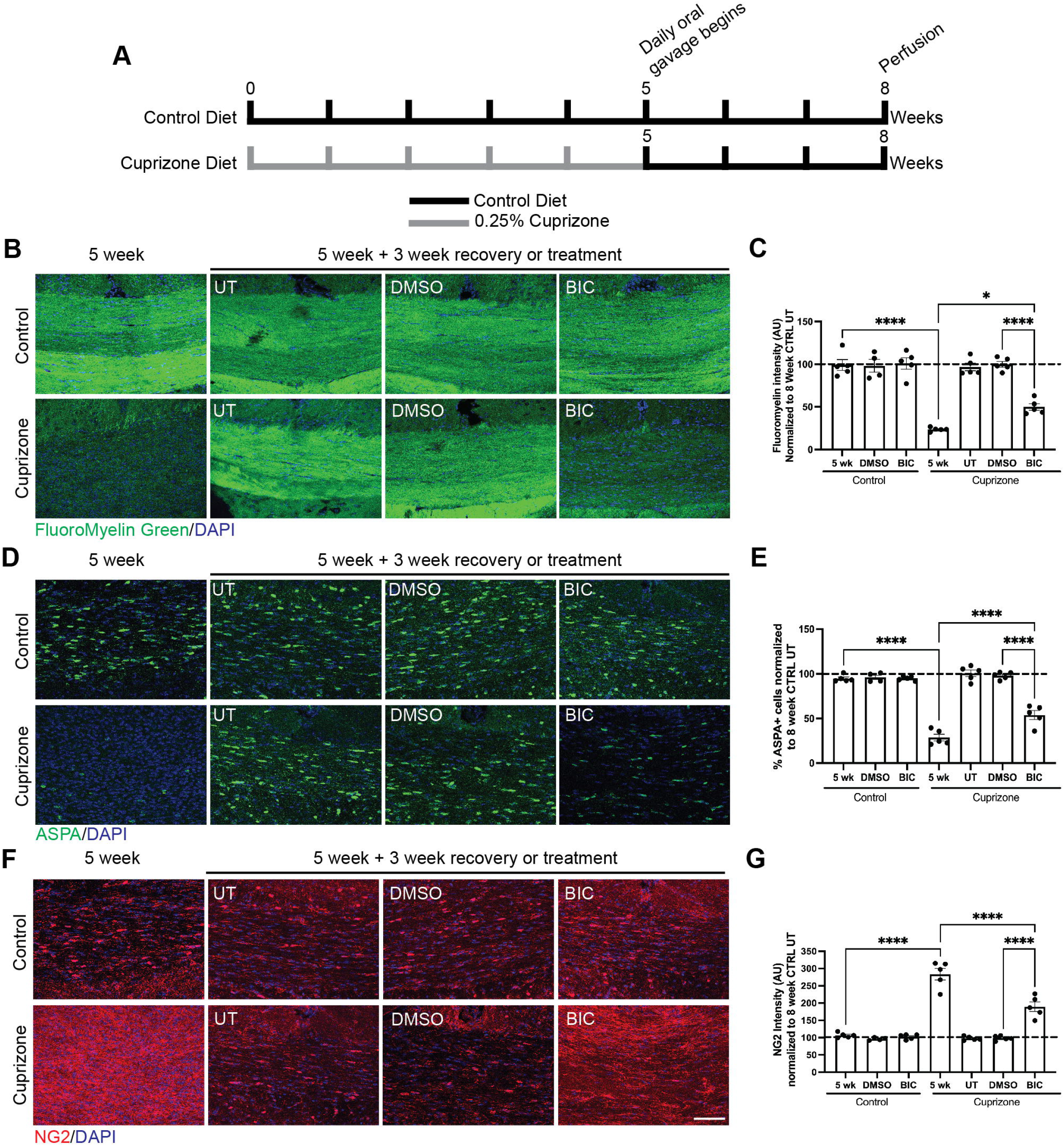
Bictegravir significantly inhibits remyelination following toxin-mediated demyelination in the corpus callosum. (A) Schematic depicting the cuprizone-mediated demyelination experiment. (B) Representative images of FluoroMyelin Green staining in the corpus callosum of control and cuprizone-fed animals treated with DMSO or BIC. (C) A significant attenuation in remyelination, as observed via a reduction in FluoroMyelin Green staining, was seen in mice fed cuprizone for 5 weeks and treated once daily with BIC during the recovery period compared to mice treated with only DMSO; *p<0.05, ****p<0.0001. (D) Representative images of mature oligodendrocyte (ASPA+) staining in the corpus callosum of control and cuprizone-fed mice. (E) Fewer mature oligodendrocytes (ASPA+) were found after the remyelination period in BIC-treated, cuprizone fed mice compared to vehicle-treated counterparts; ****p<0.0001. (F) Representative images of OPC (NG2+) staining in the corpus callosum in control and cuprizone-fed mice. (G) There were significantly more OPCs (NG2+) after remyelination in animals treated with BIC compared to vehicle-treated counterparts, suggesting an impairment of OPC’s ability to differentiate into mature oligodendrocytes; *****p<0.0001. Scale bar = 50μm. One-way ANOVA followed by Tukey’s post-hoc test. Each data point represents a single animal and data are expressed as mean ± SEM with 4-5 animals per group. All data normalized to 8-week control-fed, untreated mice.

First, myelination in the medial corpus callosum was assessed using FluoroMyelin Green, a rapid and selective dye for myelin. The 5-week cuprizone-treated mice exhibited a robust decrease in myelin, which was completely restored during the 3-week recovery period. Interestingly, mice treated with BIC during the recovery phase had a significantly attenuated remyelination response following cuprizone feeding compared to UT and vehicle-treated mice (Figure 1B & C). BIC appeared to have no impact on mature myelin, as no reduction in FluoroMyelin Green staining was observed in control diet mice treated with BIC (Figure 1B & C), suggesting that BIC does not grossly affect preexisting myelin but instead impairs remyelination.

To further validate the changes in myelination and gain insight into changes in the cells of the oligodendrocyte lineage, we stained for aspartoacylase (ASPA) for mature oligodendrocytes and neuro/glial antigen 2 (NG2) for OPCs. In line with our FluoroMyelin data, 5-week cuprizone mice had significantly fewer ASPA+ mature oligodendrocytes compared to 5-week control mice (Figure 1 D & E). Following the 3-week recovery period, the number of ASPA+ cells in untreated cuprizone mice was comparable to control mice; however, BIC-treated cuprizone mice maintained significantly reduced numbers of ASPA+ cells compared to any control mice (Figure 1D & E). Since remyelination is primarily dependent on oligodendrocyte differentiation, we also sought to identify potential alterations in OPC numbers. As previously observed in this model, following a demyelinating insult, 5-week cuprizone mice had an increase in NG2 intensity, due to the proliferation and migration of OPCs to the site of injury (Nait-Oumesmar et al., 2007), compared to 5-week control animals; this returned to basal levels at the end of the 3-week recovery period (Figure 1F & G). Interestingly, BIC-treated mice maintained elevated NG2 intensity after recovery compared to any of the control mice, though it was still significantly less than 5-week cuprizone mice (Figure 1F & G). Taken together, these data suggest that BIC significantly impairs remyelination capacity via impairment of oligodendrocyte maturation from OPCs, but does not grossly alter white matter, oligodendrocyte number or OPC numbers.

### Bictegravir does not prolong the inflammatory response induced by cuprizone

Cuprizone feeding, in addition to demyelination, is known to induce activation of astrocytes and microglia, which eventually resolves during the recovery period (Sen et al., 2022). Our previous work with elvitegravir, a member of the same drug class as BIC, demonstrated that neuroinflammation persisted in mice treated with this drug during the recovery period; therefore, we sought to determine whether a similar phenomenon was observed in BIC-treated animals. As expected, 5-week cuprizone treatment resulted in a significant upregulation of both astrocyte (GFAP+) and microglia (Iba1+) activation that was not observed in 5-week controls. Interestingly, in contrast to what we observed previously with elvitegravir (Roth et al., 2021), BIC did not result in a prolonged neuroinflammatory response when given during the recovery period, as observed by no significant differences in GFAP+ or Iba1+ reactivity (Figure 2A-C). Thus, our *in vivo* data suggest that the effects of BIC on oligodendrocyte differentiation are independent of persistent inflammation.

**Figure 2.**
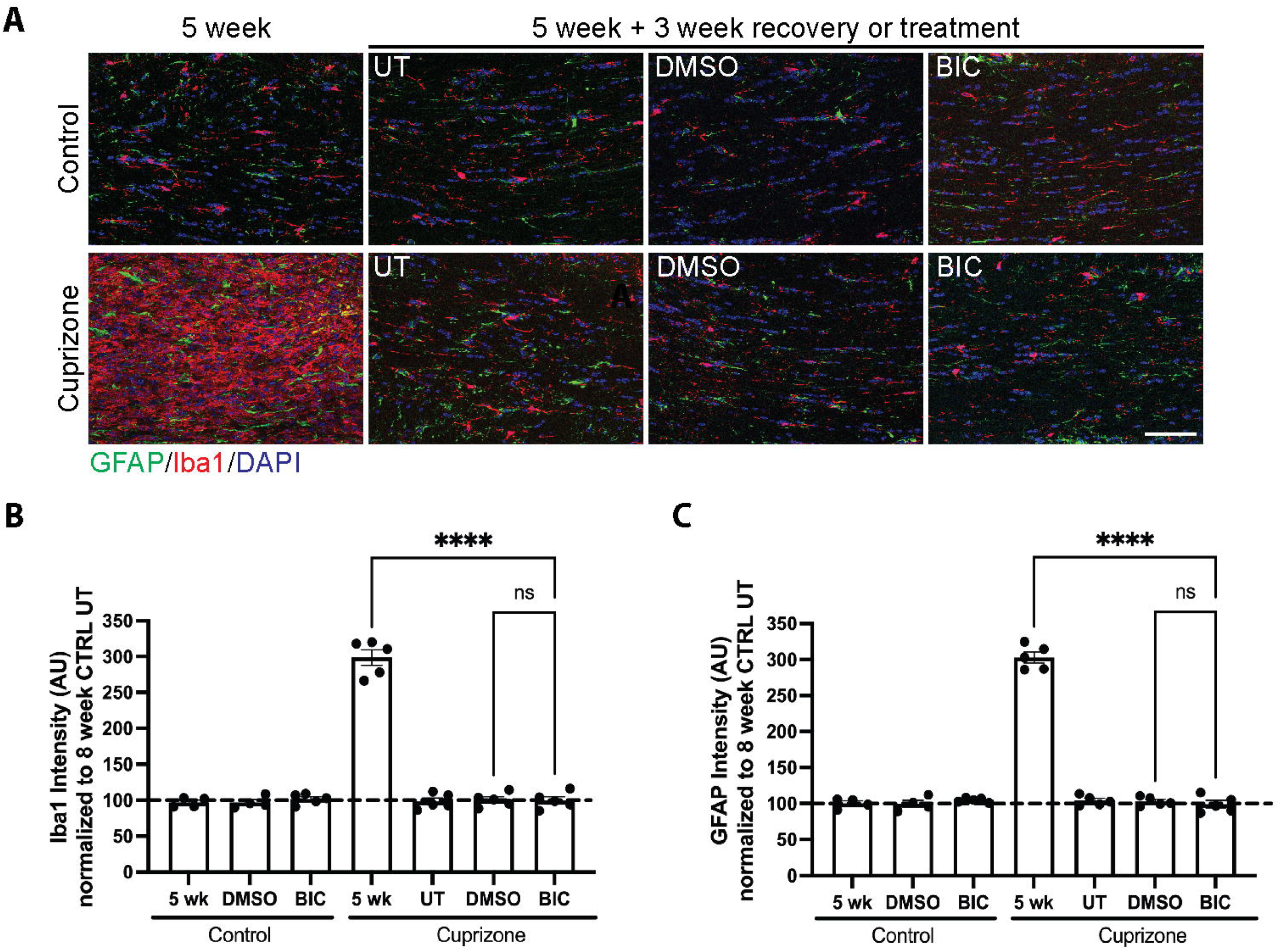
Bictegravir does not prolong the neuroinflammatory response caused by cuprizone. (A) Representative images of astrocyte (GFAP+) and microglia (Iba1+) reactivity in the corpus callosum of control and cuprizone-fed mice treated with DMSO or BIC. (B) 5 weeks of cuprizone feeding results in a significant increase in microglia reactivity that resolves following a 3-week recovery period regardless of treatment; ****p<0.0001. (C) Similar to microglia, astrocyte reactivity is significantly upregulated following cuprizone-mediated demyelination and is dampened at the end of remyelination. This is unaffected by treatment with BIC; ****p<0.0001. Scale bar = 50μm. One-way ANOVA followed by Tukey’s post-hoc test. Each data point represents a single animal and data are expressed as mean ± SEM with 4-5 animals per group. All data normalized to 8-week control diet, untreated mice.

### Bictegravir impairs oligodendrocyte differentiation in vitro

Given the observed effects of BIC on oligodendrocytes *in vivo*, we sought to determine the potential mechanisms underlying the deleterious effects of BIC on oligodendrocytes. For these *in vitro* experiments, we utilized a well-established cell culture system to model oligodendrocyte maturation (Feigenson et al., 2009) in which primary rat cortical OPCs are stimulated to differentiate in the presence or absence of BIC for 72 hours and assessed for viability and maturation. Cultures were treated with three different concentrations of BIC (1.35 μM, 13.5 μM, 41 μM), which were based on the plasma Cmax concentrations in HIV infected individuals (Gallant et al., 2017) and the number of cells expressing the following stage-specific lineage markers were quantified: A2B5 (OPCs), galactocerebroside (GalC; immature oligodendrocytes), and proteolipid protein (PLP; mature oligodendrocytes). We observed a significant decrease in GalC+ and PLP+ cells at all three concentrations of BIC after 72 hours (Figure 3B-D). To determine whether cell death or altered OPC number were responsible for these changes, we quantified the total number of DAPI positive cells and OPCs. No significant differences in OPC number or total DAPI counts were observed across all three concentrations of BIC, consistent with our previous observations that a subset of OPCs fail to proliferate in differentiation medium (Feigenson et al., 2009; Festa et al., 2021; Roth et al., 2021). Importantly, there were significantly more A2B5+ OPCs present in cultures maintained in growth media (Figure 3E-G). Since we observed a reduction in the number of PLP+ cells following BIC, we wanted to examine whether this was correlated with a decrease in myelin protein production (MBP and PLP). Interestingly, significant reductions in MBP and PLP protein levels were only observed at the highest concentration of BIC (Figure 4A-D), in contrast with our immunofluorescence data. While this finding appears to be unusual, it is not unprecedented in oligodendrocytes. Others have demonstrated that while overall protein levels of PLP and MBP can remain unchanged, localization of both proteins can be disrupted, leading to alterations in immunofluorescence counts (Maier et al., 2009; Frid et al., 2015; Jensen et al., 2015).

**Figure 3.**
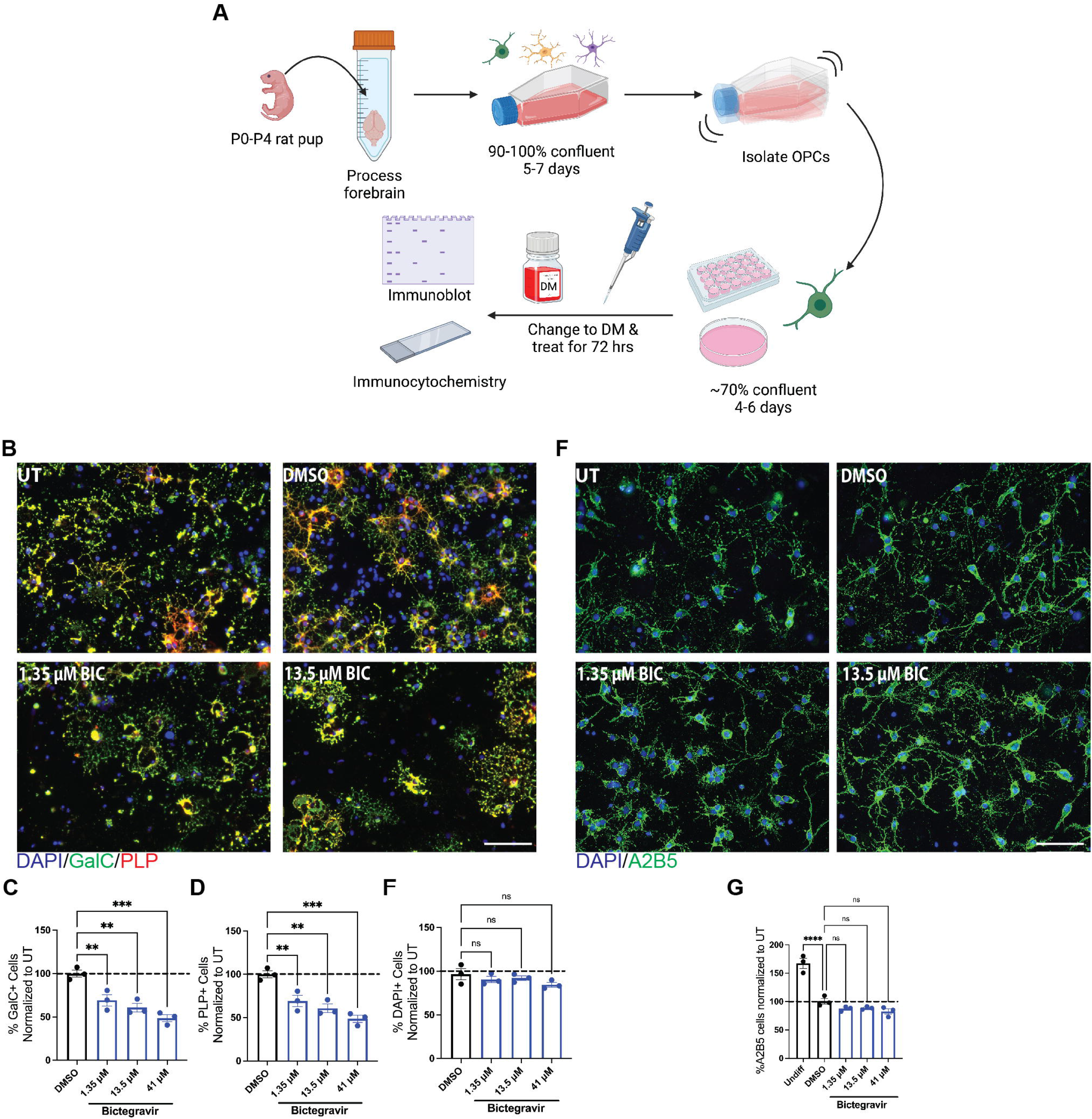
Oligodendrocyte differentiation is inhibited by bictegravir *in vitro*. (A) Schematic depicting *in vitro* treatment paradigm where purified rat OPCs are treated at the time of differentiation for 72 hours. Created with BioRender.com (B) Representative images of immature (GalC+) and mature oligodendrocyte (PLP+) staining following 72-hour differentiation in the presence of DMSO or various physiologically relevant concentrations of BIC. (C) Treatment with BIC resulted in significant decreases in the percentage of immature oligodendrocytes (GalC+) across all three concentrations tested; **p<0.01, ***p<0.001. (D) As with GalC, there was a reduction in the percentage of mature oligodendrocytes (PLP+) following treatment with BIC; **p<0.01, ***p<0.001. (E) No changes were observed in overall cell number (DAPI+) between vehicle and BIC-treated cultures. (F) Representative staining of cultures for the OPC marker A2B5 in vehicle and BIC-treated groups. (G) No significant changes in OPC (A2B5+) numbers were observed across all three concentrations of BIC; this contrasts with cells that were kept in growth media and allowed to proliferate for an additional 72 hours (Undiff). Scale bar=50 μm. One-way ANOVA followed by Tukey’s post-hoc test. Each data point depicts a biological replicate and all data normalized to and expressed as a percentage of untreated; N=3.

**Figure 4.**
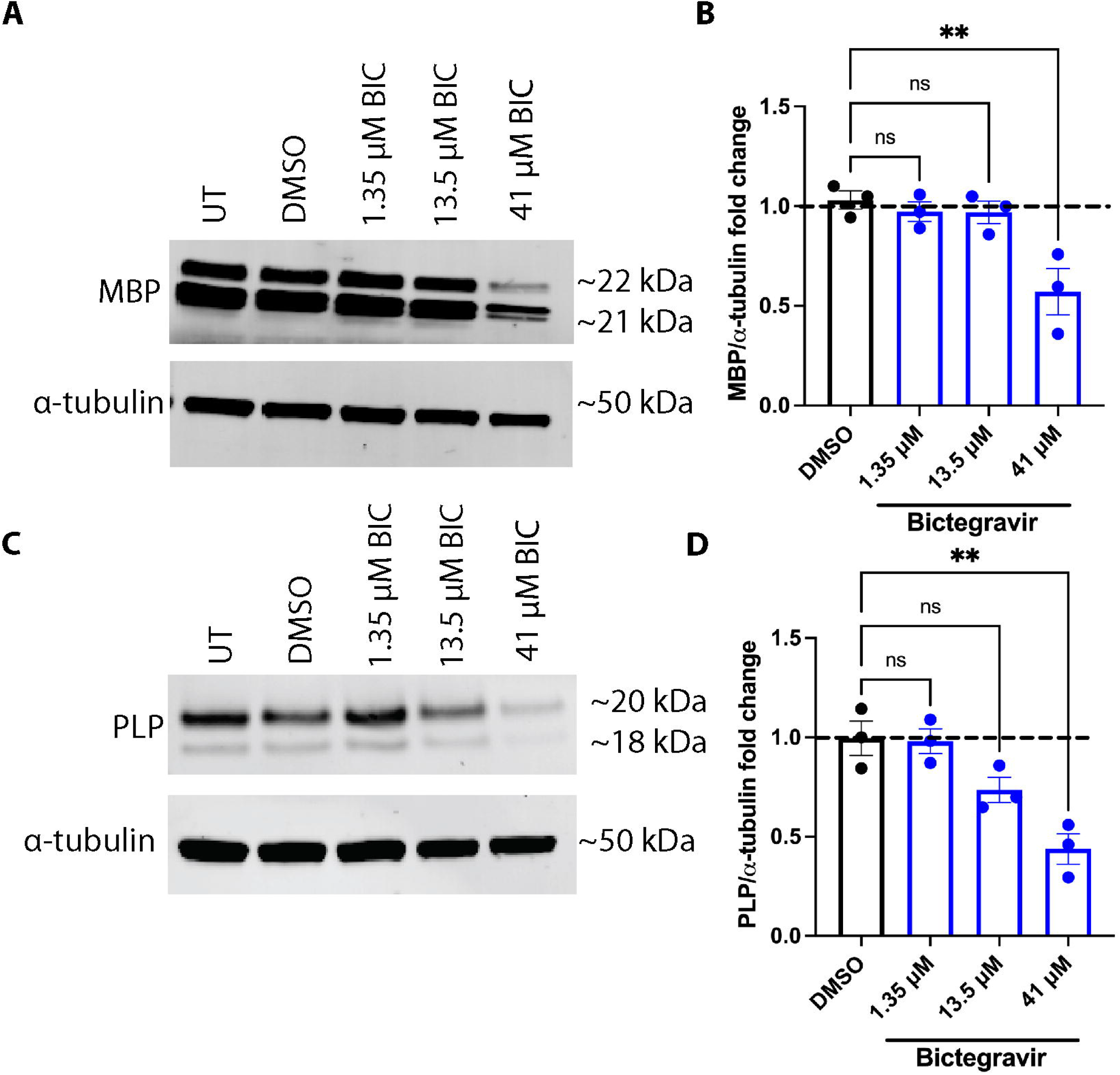
Bictegravir inhibits myelin protein production only at the highest concentration tested. (A) Representative western blots from cultures treated with vehicle or BIC for 72 hours and probed for the major myelin protein MBP and α-tubulin as a loading control. (B) A significant decrease in MBP protein was only seen at the highest concentration of BIC (41 μm) while no changes were observed at 1.35 and 13.5 μm; **p<0.01. (C) Representative western blots from vehicle and BIC-treated cultures that were assessed for the major myelin protein PLP and α-tubulin as a loading control. (D) Similar to MBP, a significant decrease in PLP was only observed at the highest concentration of BIC compared to vehicle-treated cells; **p<0.01. One-way ANOVA followed by Tukey’s post-hoc test. Each data point represents a biological replicate and all data normalized to untreated and expressed as relative fold change; N=3.

While the lack of an effect on OPCs suggests that BIC inhibits oligodendrocyte maturation and has no deleterious effects on proliferation or cell survival, we next investigated whether BIC induces OPC cell death. An increase in TUNEL+ or PI+ cells was only observed at the highest concentration of BIC (41 µM) with no significant differences seen at 1.35 µM and 13.5 µM (Figure 5A-C). Together, our data demonstrate that BIC inhibits oligodendrocyte maturation by interfering with the differentiation process rather than reducing OPC number via cell death, except for the highest concentration

**Figure 5.**
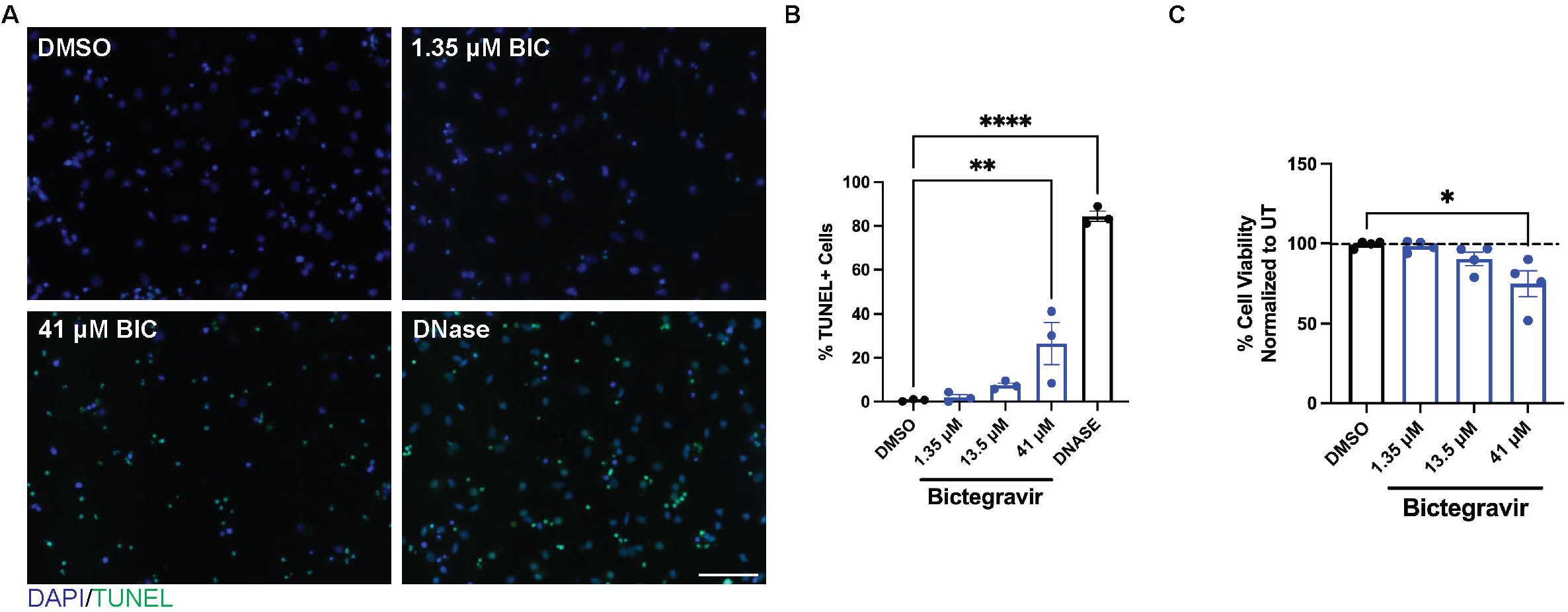
Bictegravir alters cell viability only at the highest concentration. (A) OPC cultures treated at the time of differentiation with DMSO or BIC at the time of differentiation and then assessed for apoptotic cell death via TUNEL staining. (B) Significant apoptotic cell death was only observed with 41 μM BIC after 72 hours of treatment. DNase was used as a positive control for the TUNEL assay; **p<0.01, ****p<0.0001. (C) Differentiated oligodendrocyte cultures treated with DMSO or BIC were assessed for overall cell viability using propidium iodide staining. There was a significant decrease in cell viability only at the 41 μM BIC; *p<0.05. Scale bar=50 μm. One-way ANOVA followed by Tukey’s post-doc test. Each data point represents a biological replicate and all data normalized to untreated; N=3 for TUNEL and N=4 for PI.

### Bictegravir alters lysosomal pH in OPCs and oligodendrocytes

Our previous work, along with others, has shown that other ARVs can negatively impact lysosomal function, specifically by de-acidifying the intraluminal space (Hui et al., 2019; Festa et al., 2021). The lysosome is responsible not only for recycling cellular components but has also been implicated in numerous homeostatic processes such as nutrient sensing, metabolic regulation, and endocytic sorting of major myelin proteins (Trajkovic et al., 2006; Winterstein et al., 2008; Ballabio and Bonifacino, 2020). Intriguingly, several diseases associated with lysosomal dysfunction, such as mucolipidosis type IV and Krabbe disease, are characterized by white matter loss and pallor, suggesting that the lysosome plays a critical role in modulating oligodendrocyte health (Faust et al., 2010); however their role in oligodendrocyte differentiation has not been interrogated. Thus, we sought to investigate whether BIC negatively impacts lysosomal pH.

As outlined in Figure 3A, differentiating oligodendrocytes were exposed to BIC and the number of acidic organelles were quantified by counting the number of LysoTracker Red+ puncta, a cell-permeant acidtropic dye, in GalC+ and A2B5+ cells. We observed fewer acidic organelles in both OPCs (A2B5+) and immature/mature oligodendrocytes (GalC+) in cultures treated with BIC, suggesting that BIC may disrupt lysosomal acidity (Figure 6A-D). While LysoTracker Red staining provides information about the number of acidic organelles present within a cell, it fails to inform us about the quantitative pH changes that are occurring. To investigate this, we utilized the ratiometric probe, LysoSensor Yellow/Blue DND-160, to measure the pH of acidic organelles, including lysosomes, which typically have an optimal pH of 4.5-5.0. Following 72-hour treatment with BIC, we observed significant de-acidification of acidic organelles at 13.5 μM and 41 μM with a mean pH of 5.415 and 5.539 respectively (Figure 6E). Interestingly, there was no significant alkalinization of acidic organelles at the 1.35 μM concentration (mean pH of 4.686), despite a reduction in LysoTracker Red+ puncta at this concentration.

**Figure 6.**
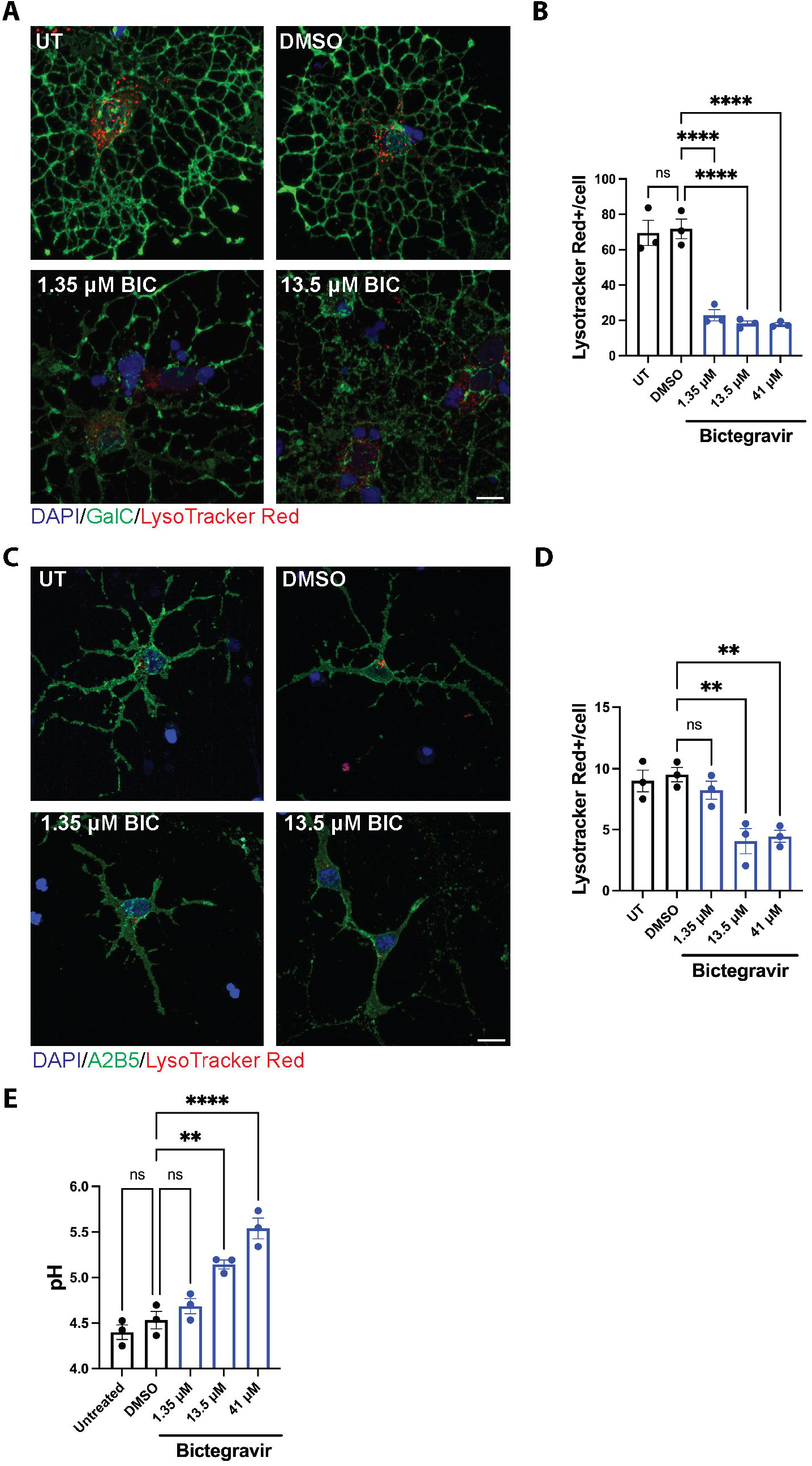
Bictegravir alters lysosomal pH in OPCs and oligodendrocytes. (A) Representative images of cells treated at the time of differentiation with DMSO or BIC and incubated with LysoTracker Red for 30 mins and stained for GalC. (B) The number of LysoTracker Red puncta per GalC+ cell was significantly decreased across all three concentrations of BIC compared to vehicle-treated cultures; ****p<0.0001. (C) OPC cultures were treated at the time of differentiation for 72 hrs, incubated with LysoTracker Red, and then stained for A2B5. (D) The number of acidic organelles was significantly decreased in the 13.5 and 41 μm BIC treatment groups in OPCs (A2B5+) compared to vehicle-treated cells; **p<0.01. (E) Lysosomal pH was assessed using LysoSensor Yellow/Blue following treatment with DMSO or BIC for 72 hours. Significant lysosomal de-acidification occurred in the 13.5 and 41 μm BIC treatment groups while no change was observed in the 1.35 μm group; **p<0.01, ****p<0.0001. Scale bar=10 μm. One-way ANOVA followed by Tukey’s post-hoc test. Each data point represents a biological replicate; N=3.

### Bictegravir impairs lysosomal degradative capacity in both OPCs and oligodendrocytes

While we have demonstrated that lysosomal pH in the OL lineage is disrupted upon BIC treatment, the above-reported assays do not provide insight into the functionality of these lysosomes. Therefore, we assessed lysosomal proteolytic activity using DQ-BSA Red as our functional readout. In this assay, BSA is heavily labeled with BODIPY TR-X dye, resulting in self-quenching, and is loaded into cells for 6 hours. This cargo is endocytosed and trafficked through early endosomes to late endosomes, which eventually fuse with acidic lysosomes. In these endolysosomes, DQ-BSA Red is broken down into smaller fragments, which leads to dequenching of the fluorophores and bright fluorescence within acidic compartments. Seventy two-hour treatment with BIC during differentiation resulted in significantly fewer DQ-BSA Red+ puncta in OPCs and OLs (Figure 7A-D), indicating a reduction in the number of functional, degradative organelles. Taken together, these data demonstrate that physiologically relevant concentrations of BIC result in lysosomal de-acidification and impaired lysosomal proteolytic activity in both OPCs and oligodendrocytes, thus providing a potential mechanism by which BIC inhibits oligodendrocyte differentiation.

**Figure 7.**
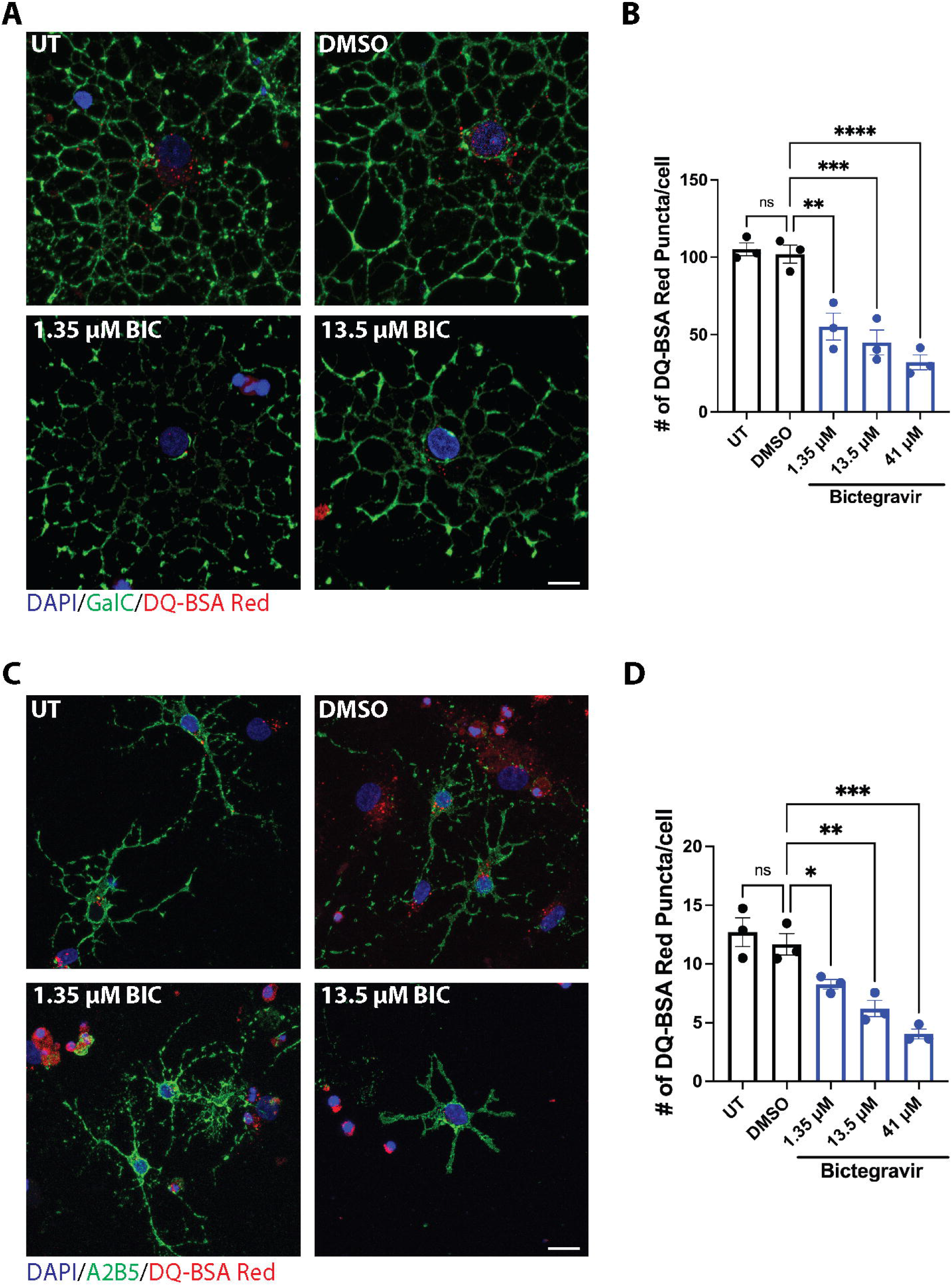
Bictegravir impairs lysosomal proteolytic activity in OPCs and oligodendrocytes. (A) Differentiated oligodendrocytes treated with either DMSO or BIC were incubated with DQ-BSA Red for 6 hours and then stained for GalC. (B) The number of proteolytically active lysosomes were significantly decreased in immature/mature oligodendrocytes following treatment with BIC; **p<0.01, ***p<0.001, ****p<0.0001. (C) Cells treated with DMSO or BIC were incubated with DQ-BSA Red for 6 hours and stained for the OPC marker A2B5. (D) There were fewer proteolytically active lysosomes in OPCs (A2B5+) treated with BIC compared to vehicle-treated cultures; *p<0.05, **p<0.01, ***p<0.001. Scale bar=10 μm. One-way ANOVA followed by Tukey’s post-hoc test. Each data points represents a biological replicate; N=3.

### Bictegravir does not alter lysosomal membrane integrity or lysosome number

While our data thus far is indicative of lysosomal acidification defects being a primary driver of BIC-mediated inhibition of oligodendrocyte maturation, it does not rule out a reduction in overall lysosome number. To address this, we first examined whether lysosomal membrane integrity was affected by BIC treatment. Galectin-3 belongs to a family of cytosolically synthesized lectins with an affinity for β-galactoside glycoconjugates that are typically present on the luminal membrane of several organelles, including lysosomes (Johannes et al., 2018). Upon lysosomal membrane disruption, galectin-3 is recruited to lysosomes where it forms discernible puncta as assessed via ICC (Aits et al., 2015). Following BIC exposure during the 72-hour differentiation period, we did not observe an increase in Gal-3+ puncta across any of the three concentrations used; in contrast, a 2 hr treatment with the lysosomal-membrane-damaging agent Leu-Leu-OMe (LLOME; 1 mM) resulted in a significant increase in the number of Gal3+ puncta around DAPI+ nuclei (Figure 8A-B). Therefore, BIC does not seem to induce lysosomal membrane disruption that could explain the de-acidification defects that we observe.

**Figure 8.**
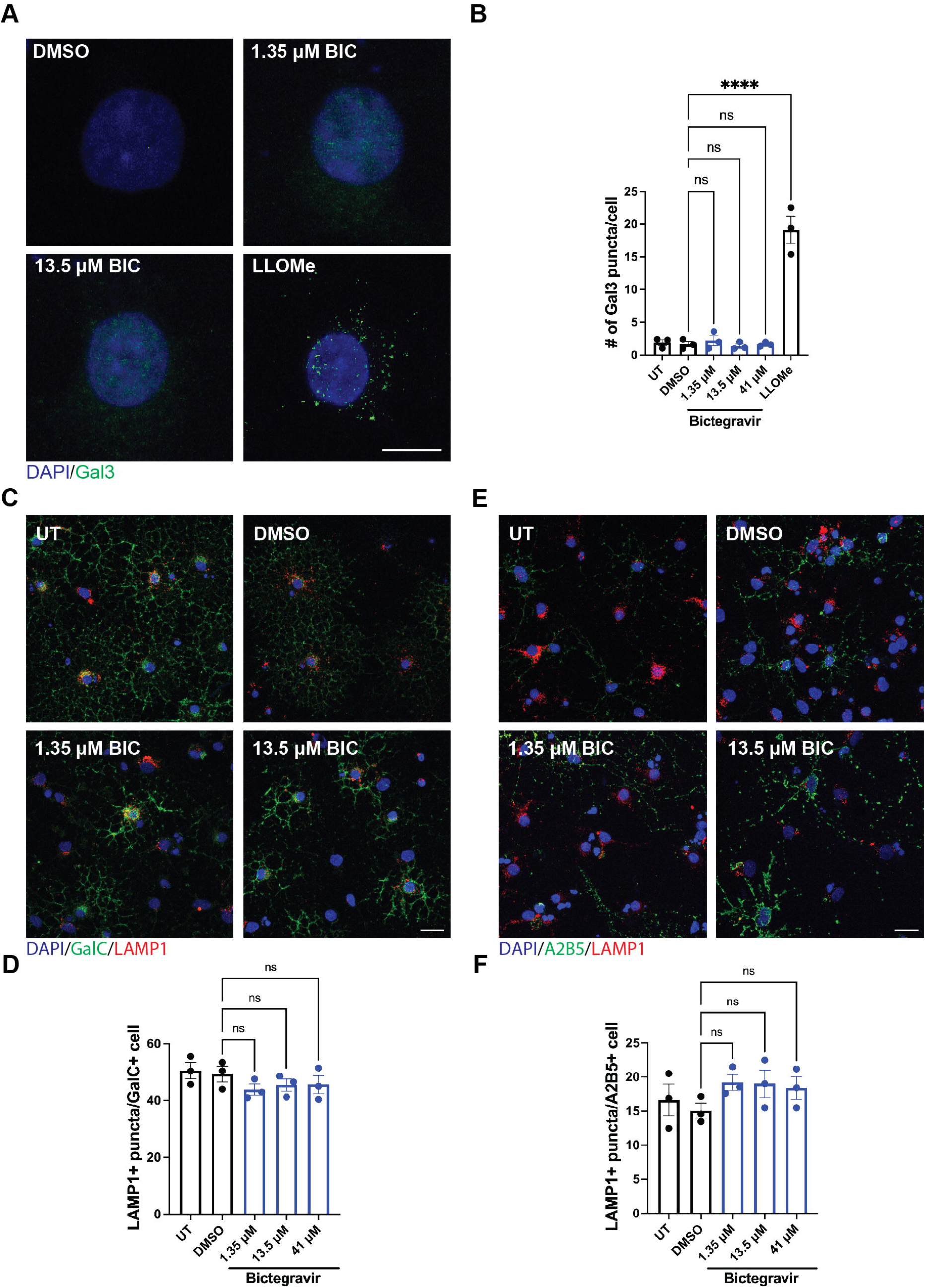
Lysosomal membrane permeability and total lysosome number are not altered by bictegravir. (A) OPC cultures were treated with DMSO or BIC at the time of differentiation for 72 hrs and then stained for galectin 3 (Gal3), a marker of lysosomal membrane damage. (B) The number of Gal3+ puncta in BIC-treated cultures were not significantly different from untreated or vehicle-treated cells. Two-hour treatment with LLOMe resulted in a large number of perinuclear localized Gal3+ puncta. (C) Following 72 hrs of DMSO or BIC treatment, cells were stained for the lysosomal membrane protein LAMP1 and the immature/mature oligodendrocyte marker GalC. (D) Treatment with BIC did not alter the overall number of lysosomes in GalC+ cells. (E) In a similar manner, cells treated with DMSO or BIC were stained for LAMP1 and the OPC marker A2B5. (F) As with immature/mature oligodendrocytes, BIC treatment did not change the total number of lysosomes in OPCs (A2B5+). Scale bar=10μm. One-way ANOVA followed by Tukey’s post-hoc test. Each data point represents a single biological replicate; N=3.

While it appears that BIC does not cause disruption to the lysosomal membrane, it is possible that BIC could alter overall lysosome number via other mechanisms that would explain the decrease in acidic and degradative organelles. Intriguingly, total lysosome number, as assessed via staining of the lysosomal membrane glycoprotein LAMP1, was not changed following treatment with any concentration of BIC used (Figure 8C-F). Thus, our data demonstrate that BIC specifically impairs lysosomal acidification and degradative capacity without affecting lysosomal membrane integrity or overall lysosome number, which results in oligodendrocyte maturation deficits.

### De-acidification of lysosomes is sufficient to inhibit oligodendrocyte differentiation

To determine whether lysosomal de-acidification is sufficient to inhibit oligodendrocyte maturation, we treated OPCs with the vacuolar H^+^ ATPase inhibitor bafilomycin A (Baf A), which results in increased lysosomal pH, during differentiation. We first confirmed that the concentrations of Baf A utilized resulted in measurable changes in lysosome pH over the differentiation period (Figure 9A). As in our BIC experiments, differentiating OPC cultures were treated with Baf A and then stained for A2B5, GalC, and PLP to determine rates of oligodendrocyte maturation. Lysosomal de-acidification via Baf A treatment was sufficient to inhibit OPC differentiation, as we observed significantly fewer immature and mature oligodendrocytes, with no associated changes in overall cell number (Figure 9B-E). Interestingly, we also did observe a reduction in OPC number at 500 and 750 pM BafA (Figure 9F-G). Therefore, our data demonstrate that proper lysosomal acidification is critical for OL maturation to occur, and this has implications for neurologic disorders that are characterized by lysosomal dysfunction.

**Figure 9.**
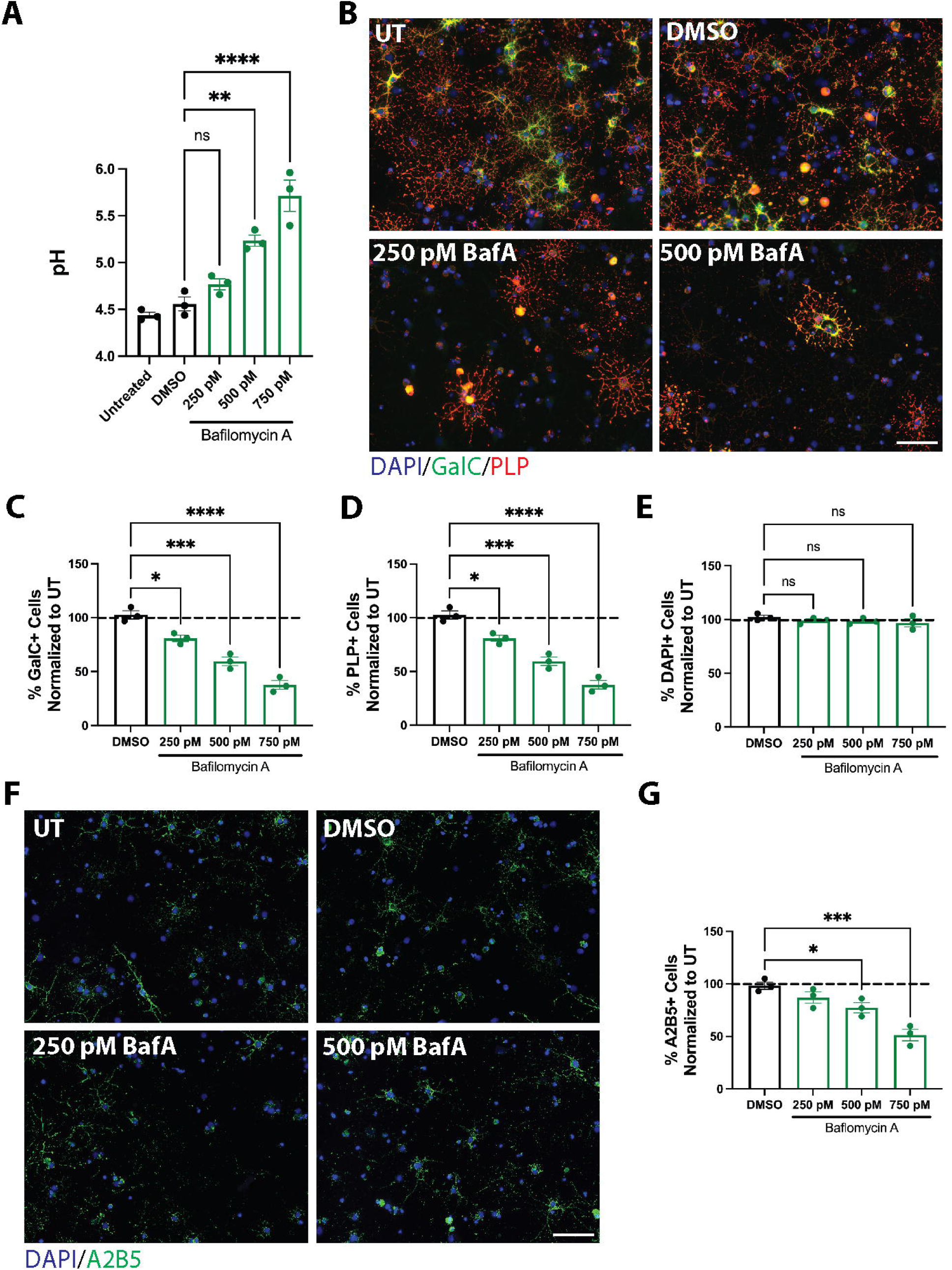
Lysosomal de-acidification is sufficient to prevent oligodendrocyte maturation. (A) OPC cultures were treated with bafilomycin A at the time of differentiation for 72 hours and then lysosomal pH was measured using LysoSensor Yellow/Blue. There was a stepwise increase in lysosomal pH as the concentration of bafilomycin A increased; **p<0.01, ****p<0.0001. (B) Cultures were treated with various concentrations of bafilomycin A during differentiation and stained for GalC and PLP as immature and mature oligodendrocyte markers, respectively. (C) The percentage of GalC+ (immature oligodendrocytes) was significantly decreased in all three concentrations of bafilomycin A tested; *p<0.05, ***p<0.001, ****p<0.0001. (D) Similarly, the percentage of mature oligodendrocytes (PLP+) were significantly decreased across all three concentrations of bafilomycin A; *p<0.05, ***p<0.001, ****p<0.0001. (E) No changes were observed in overall cell number (DAPI+) at any concentration of bafilomycin A. (F) OPC cultures were treated with bafilomycin A at the time of differentiation for 72 hrs and stained for the OPC marker A2B5. (G) There was a significant decrease in the percentage of OPCs (A2B5+) at 500 and 750 pM bafilomycin A compared to vehicle-treated cultures. Scale bar= 50 μM. One-way ANOVA followed by Tukey’s post-hoc test. Each data point represents a biological replicate and all data normalized to untreated; N=3.

### Activation of the lysosomal cation channel TRPML1 rescues both bictegravir and bafilomycin A-mediated inhibition of oligodendrocyte maturation

Lysosomal acidification is achieved by the activity of the vacuolar-type H^+^ ATPase which uses energy derived from ATP hydrolysis to transport protons across the lysosomal membrane (Forgac, 2007). This leaves an electrical potential, that if left uncompensated, can limit the ability of the V-ATPase to continue pumping and reach a sufficiently acidic pH. In order to alleviate this restraint, counter ion pathways involving the influx of anions and/or efflux of cations are necessary to dissipate the development of a restrictive electrical gradient. While there are well-established candidate transporters for the anion-moving portion of the lysosomal counterion pathway, candidates for the cation-transporting aspect are still tentative. One likely candidate cation channel is the transient receptor potential mucolipin 1 (TRPML1), which is primarily permeable to calcium, in addition to iron and zinc. Our group and others have previously shown that activation of TRPML1 is sufficient to restore de-acidification deficits in oligodendrocytes and neurons caused by antiretrovirals (Hui et al., 2019; Festa et al., 2021).

First, we demonstrated that co-administration of the TRPML1 agonist, MLSA1, prevented lysosomal de-acidification caused by BIC (Figure 10A). Once we established the effect of MLSA1 on prevent lysosomal de-acidification, we assessed whether co-treatment with MLSA1 in the presence of BIC would prevent inhibition of oligodendrocyte maturation. Activation of TRPML1 prevented BIC-induced deficits in oligodendrocyte differentiation; the levels of immature (GalC+) and mature oligodendrocytes (PLP+) were comparable to vehicle-treated levels at the low and medium concentrations of BIC. At the highest concentrations of BIC, there were significantly more oligodendrocytes; however, this did not reach baseline levels of differentiation (Figure 10B-F). Taken together, these data highlight the critical role lysosomal Ca^2+^ efflux plays in restoring lysosomal acidification during lysosomal stress. Additionally, these data may suggest a physiological role for lysosomal Ca^2+^ in modulating oligodendrocyte maturation.

**Figure 10.**
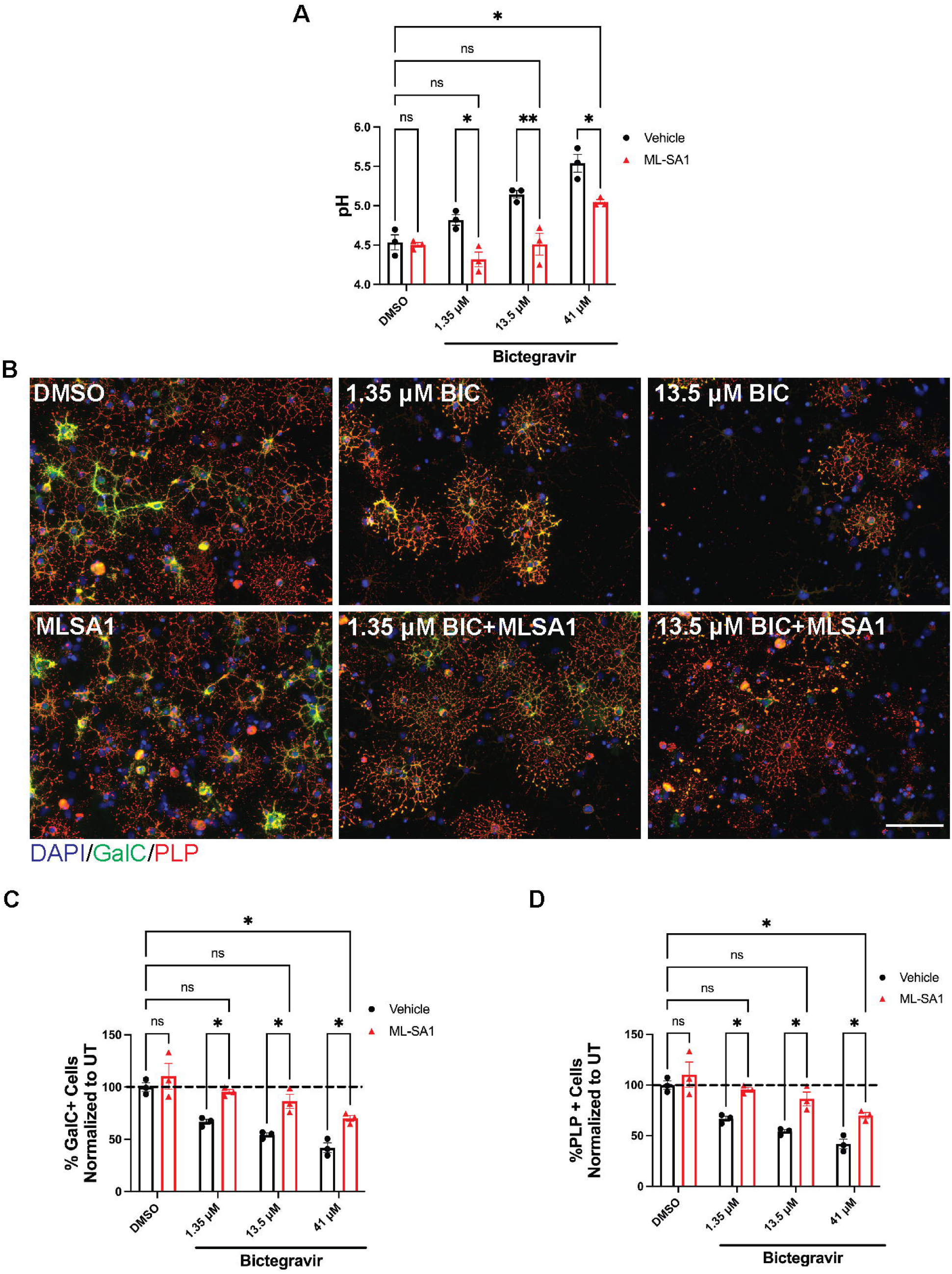
Activation of the lysosomal cation channel TRPML1 prevents bictegravir-mediated lysosomal de-acidification and inhibiting of oligodendrocyte maturation. (A) Co-administration of the TRPML1 agonist, MLSA1, prevented lysosomal de-acidification induced by BIC at the 1.35 and 13.5 μM concentration; reacidification also occurred at 41 μM BIC but it did not return to baseline pH; *p<0.05, **p<0.01. (B) OPC cultures were treated at the time of differentiation with DMSO or BIC with or without MLSA1 and then stained for GalC and PLP. (C) Co-treatment with MLSA1 prevented a decrease in the percentage of GalC+ cells at the 1.35 and 13.5 μM BIC; there was also an effect at the 41 μM BIC but this did not return to vehicle treated levels; *p<0.05. (D) Similarly, the percentage of PLP+ cells at the 1.35 and 13.5 μM BIC returned to vehicle-treated levels when cultures were co-treated with MLSA1; *p<0.05. Scale bar=50 μm. Two-way ANOVA followed by Tukey’s multiple comparisons test. Each data point represents a biological replicate and all data normalized to untreated. N=3.

## Discussion

The study is the first to demonstrate that proper lysosomal acidification is a critical determinant of oligodendrocyte maturation. We showed that the ARV drug, bictegravir, inhibits remyelination mediated via OPC differentiation without prolonging the neuroinflammatory environment induced by cuprizone-mediated demyelination. Our mechanistic studies revealed that disruption of lysosomal acidity, via BIC or other drugs (e.g. bafilomycin A) is sufficient to prevent oligodendrocyte maturation and this effect can be prevented by activation of the cation-permeable lysosomal channel, TRPML1. Thus, we have uncovered a novel mechanism regulating maturation of oligodendrocytes that has implications not only for HIV-associated neurocognitive disorders, but also other diseases characterized by white matter impairment and lysosomal dysfunction.

Remyelination is the most effective and robust endogenous repair mechanism in the CNS, which can lead to the complete recovery of large areas of demyelination in the CNS with resultant functional rescue (Lubetzki et al., 2020). While recent studies from rodent and large animal models suggest that oligodendrocytes that survive myelin breakdown can generate new sheaths and contribute to myelin pattern preservation following demyelination, the primary contributor to remyelination appears to be the migration, proliferation, and differentiation of OPCs (Bacmeister et al., 2020; Orthmann-Murphy et al., 2020). Therefore, animal models used to induce demyelination, such as cuprizone-feeding, can also provide insight on the mechanisms modulating OPC differentiation and maturation (Zirngibl et al., 2022). Since our previous work implicated another ARV drug, elvitegravir, in disruption of remyelination following cuprizone-mediated demyelination, we aimed to examine whether this was true for BIC. We found that administration of BIC during the recovery period significantly attenuated the ability of mature myelin to reform (Fig. 1B & C), consistent to what was observed with elvitegravir (Roth et al., 2021), although elvitegravir appeared to impact microglia and astrocytes as well as oligodendrocytes while the effects of BIC appear limited to the differentiating oligodendrocytes. Further analysis revealed that this deficit in myelin was driven by a failure to repopulate myelinating oligodendrocytes, as observed by significantly fewer ASPA+ cells in the corpus collosum in cuprizone-fed, BIC-treated animals (Fig 1D & E). This decrease in ASPA+ cell count appeared to be driven by inhibition of OPC differentiation as the number of OPCs (NG2+ cells) in cuprizone-fed, BIC-treated animals were significantly elevated compared to DMSO-treated counterparts (Fig 1F &G). Thus, it appears that BIC does not impact the migration of OPCs to the site of demyelination or OPC proliferation, but instead alters the ability of OPCs to mature into myelinating oligodendrocytes.

While our data demonstrates that mature oligodendrocytes fail to reemerge after demyelination in the presence of BIC, it does not rule out the possibility that newly formed, premyelinating oligodendrocytes have appeared. Premyelinating oligodendrocytes (preOLs), identified by expression of breast carcinoma amplified sequence 1 (BCAS1) and Ectonucleotide Pyrophosphatase/Phosphodiesterase 6 (ENPP6), constitute a population of terminally differentiated cells that are not OPCs but have yet to start generating myelin sheaths. Typically, a majority of these preOLs are lost, most likely due to apoptosis; those that survive this checkpoint undergo dramatic changes in gene expression and morphology that enable them to integrate into mature, myelinating oligodendrocytes (Barres and Raff, 1994; Hughes and Stockton, 2021). These changes require a high energy demand and metabolic stress; accumulating evidence indicates that preOLs are considerably more vulnerable to injury via hypoxia, limited metabolites, active microglia, and neuronal activity compared to OPCs and mature oligodendrocytes (Hughes and Stockton, 2021). It is possible that BIC exposure during a period of remyelination alters preOL survival and/or integration, resulting in fewer myelinating OLs and inhibition of functional recovery. On the other hand, the accumulation of OPCs may be indicative of a failure for these cells to emerge from this early stage of the lineage. Surprisingly, in contrast to what we previously observed with elvitegravir, animals treated with BIC after cuprizone-mediated demyelination had partial remyelination occur during the 3-week recovery period (Figure 1C & E). This raises the possibility that this observation is due to a delay in remyelination caused by BIC or reflects a heterogeneous OPC response to injury. Previous work with elvitegravir, *in vitro*, demonstrated that upon washout of the drug, differentiation was able to proceed normally for one of the concentrations tested (Roth et al., 2021). Thus, it seems possible that BIC-treated, cuprizone fed animals would remyelinate following cessation of BIC treatment. Heterogeneity of OPCs may also contribute to the partial remyelination we observed following exposure to BIC. It is now well-established that there are a subset of OPCs that express the G-protein coupled receptor 17 (GPR17), a protein that is expressed during the progenitor and premyelinating phase and is downregulated in order for maturation to occur (Nyamoya et al., 2019). In response to a demyelinating injury, such as cuprizone, the pool of GPR17+ OPCs rapidly expands and accumulates in the lesions; during the recovery period, these cells are able to successfully proceed to terminal differentiation and express myelin proteins (Coppolino et al., 2018). However, there have been reports that sustained upregulation of GPR17 results in remyelination failure, though the exact mechanism that underlies this continued expression is unknown (Coppolino et al., 2018; Wang et al., 2020). The few OPCs that were able to differentiate into mature oligodendrocytes even in the presence of BIC were possibly GPR17-negative, while the remaining OPCs continuously expressed high levels of GPR17; ongoing work in the laboratory is investigating the contribution of GPR17 to remyelination failure in the context of ARVs.

One of the most surprising findings from our *in vivo* studies was that exposure to BIC during remyelination did not prolong the neuroinflammatory response, as measured by astrocytic (GFAP+) and microglial (Iba1+) reactivity, induced by cuprizone (Figure 2A-C). This is in stark contrast to what was observed with elvitegravir, where activation of both astrocytes and microglia were observed at the end of the recovery period (Roth et al., 2021). Recruitment of both of these cells, as well as their ability to phagocytose myelin debris, are essential for remyelination to occur after cuprizone-mediated demyelination; however, their prolonged activation can lead to the secretion of deleterious substances that impede remyelination (Sen et al., 2022). It is possible that BIC is impacting the ability of astrocytes and microglia to phagocytose myelin debris or altering their secretome, which in turn impair myelin regeneration. Intriguingly, recent work has demonstrated the neuroinflammatory reactive astrocytes undergo lysosomal remodeling and exocytosis that underlies astrocyte neurotoxicity (Rooney et al., 2021).

Our mechanistic *in vitro* studies revealed lysosomal de-acidification as a significant driver of BIC-mediated inhibition of oligodendrocyte maturation (Figures 5-8). Furthermore, proper lysosomal acidification is necessary in order for OPCs to differentiate into mature oligodendrocytes (Figure 9). Recently, there has been burgeoning interest in understanding the complex functions of the lysosome outside of its traditional role as the primary degradative organelle of the cell. In oligodendrocytes, little research has been performed on the specialized role of the lysosome in regulating oligodendrocyte health and function. The transport of the major myelin protein, PLP, is known to be dependent on lysosomal exocytosis; furthermore, the myelin proteins, MAG and MOG, as well as the OPC proteoglycan, NG2, depends on endocytic recycling through the lysosome (Trajkovic et al., 2006; Winterstein et al., 2008; Daynac et al., 2018). However, the impact of lysosomal acidification on these processes has not been assessed. Folts et al (2016) demonstrated that structurally related sphingolipids known to accumulate in lysosomal storage disorders (LSDs) were sufficient to disrupt lysosomal pH in OPCs and this resulted in deficits in OPC proliferation and survival (Folts et al., 2016). Our data now demonstrates that proper lysosomal acidification is indispensable for OPCs to terminally differentiate into mature oligodendrocytes. We observed that treating differentiating OPCs with either BIC or Baf A resulted in significant inhibition of immature and mature oligodendrocyte formation (Figures 3 & 9), and this was associated with significant lysosomal alkalinization (Figures 6 & 9) and impaired lysosomal degradative capacity (Figure 7). While our data has revealed the critical role of lysosomal acidification in oligodendrocyte differentiation, the exact processes that are disrupted by lysosomal de-acidification are still unknown.

Unexpectedly, while we found significantly fewer PLP+ mature oligodendrocytes following BIC treatment (Figure 3A & D), there were only significant changes in PLP protein expression at the highest concentration (Figure 4C & D), most likely a reflection of cell death at this concentration (Figure 5). Since this phenomenon, where myelin protein expression remains unchanged but their localization is altered, has been observed before (Maier et al., 2009; Frid et al., 2015; Jensen et al., 2015), it seems likely that proper localization of PLP via lysosomal exocytosis is pH dependent. Therefore, it is possible that lysosomal de-acidification in maturing oligodendrocytes leads to differentiation defects by blocking the insertion of PLP into the plasma membrane, where the protein, instead, accumulates with lysosomal compartments; however, this requires further investigation. Additionally, this potential accumulation of PLP within the lysosome may not only impact the protein composition of the myelin membrane but may also impact the lipid content. The myelin sheath is characterized by a high proportion of lipids (70-85%) compared to proteins (15-30%), which contributes to the tight packing and organization of myelin through non-covalent interactions between lipids and myelin proteins (Min et al., 2009). Cholesterol is one of the most abundant lipids present in the myelin sheath (∼40%) and is known to physically interact with PLP within the lysosome (O’brien, 1965; Simons et al., 2000). Studies in a Pelizaeus-Merzbacher disease (PMD) mouse model demonstrated that accumulation of PLP within lysosomes leads to increased intracellular cholesterol content and subsequent inhibition of myelin assembly due to insufficient cholesterol in the plasma membrane (Saher et al., 2012).

While our *in vitro* work focused on the differentiation capacity of oligodendrocytes following lysosomal de-acidification, this does not address how it may impact already mature, myelinating oligodendrocytes. In animals fed control diet for the entire 8 weeks and treated with BIC for the last 3 weeks, we observed no gross changes in mature myelin or the number of mature oligodendrocytes (Figure 1A-D), suggesting that mature oligodendrocytes might be resistant to the effects of lysosomal de-acidification; however, it is important to note that these animals were treated with BIC for a relatively short period of time and people living with HIV are typically taking these medications for many years. This is important in light of recent findings demonstrating that, unlike previously thought, mature myelin is metabolically active and undergoes rapid turnover of two major lipid components, phospholipids and sphingolipids (Zhou et al., 2020). Since lysosomes are responsible for the process and sorting of lipids, as well as metabolic control hub, it seems feasible that long-term treatment with BIC or other ARVs that induce lysosomal de-acidification may impact the lipid turnover that is necessary to maintain myelin and lead to white matter dysfunction observed in these individuals. In support of this, the duration of ART treatment is correlated with the amount of white matter attenuation in people living with HIV and myelin gene transcripts are known to remain dysregulated in individuals taking ART (Borjabad et al., 2011; Seider et al., 2016; Solomon et al., 2019).

TRPML1 is non-selective cation permeable channel that is localized to late endosomal/lysosomal compartments, where it functions to extrude primarily Ca^2+^ from the lysosomal lumen and into the cytosol. This channel is known to play essential roles in lysosomal biogenesis, exocytosis, and positioning, as well as intraorganellar Ca^2+^ signaling with the endoplasmic reticulum and mitochondria (Di Paola et al., 2018; Griffin et al., 2020; Peng et al., 2020). In addition to these above-mentioned functions, TRPML1 has emerged as a candidate counterion efflux channel to dissipate the buildup of a restrictive electrical gradient generated by the V-ATPase. Our current data demonstrating that activation of TRPML1 can prevent BIC-induced lysosomal de-acidification from occurring in oligodendrocytes (Figure 10), as well as data from others demonstrating a similar effect in neurons, provides compelling evidence that Ca^2+^ efflux is important for regulating the electrical gradient necessary to maintain proper V-ATPase function (Bae et al., 2014; Hui et al., 2019; Festa et al., 2021). Additionally, these data suggest that lysosomal Ca^2+^ may play an essential role in regulating oligodendrocyte maturation. In line with this, inactivating mutations in TRPML1 leads to the rare disease mucolipidosis type IV, which is characterized by white matter abnormalities and a hypoplastic corpus callosum (Misko et al., 2021). Work done in *Mcoln1*^*-/-*^ (the gene that encodes TRPML1) mice has helped elucidate the nature of myelination deficits resulting from inactivation or loss of TRPML1, including reduction in a number of mature oligodendrocyte markers that were not due to a loss of OPCs or neuronal fibers (Grishchuk et al., 2015; Mepyans et al., 2020). These data, along with our findings regarding the importance of TRPML1 in alleviating lysosomal de-acidification, implicates a critical role of TRPML1 in modulating oligodendrocyte function; however, the exact mechanism underlying these observations is unknown.

Here, we highlight a previously unappreciated role of the lysosome in modulating oligodendrocyte maturation. Additionally, we provide further evidence that modulation of the endolysosomal cation channel, TRPML1, is sufficient to alleviate both lysosomal de-acidification and oligodendrocyte maturation deficits induced by ARVs. This has implications not only for people with HIV on suppressive ART but also for other neurologic disorders where lysosomal acidification defects are starting to be unveiled, including Alzheimer’s disease and Parkinson’s disease. Thus, therapeutic targeting of either TRPML1 or lysosomal acidification may represent a viable strategy to alleviate white dysfunction across a spectrum of neurological diseases.

## Acknowledgments

This project was supported by the following grants: R01 MH098742 (KJS and JBG), R01 MH126773 (KJS and JBG), R21 MH118121 (KJS and JBG) and T32 AI007632 to the Department of Microbiology at the University of Pennsylvania (LKF). We also thank the laboratory of Michael Robinson, PhD at the Children’s Hospital of Philadelphia for the use of the Odyssey Infrared Imaging System and the Leica DMi8 confocal microscope, and the Mass Spectrometry Core at the Children’s Hospital of Philadelphia. The authors declare no competing financial interests. Table of contents image created with Biorender.com.

## Notes

**Data availability statement** The data that support the findings of this study are available from the corresponding authors, Kelly Jordan-Sciutto and Judith Grinpsan, upon request.

### Competing Interest Statement

The authors have declared no competing interest.

